# Assessing the impact of successive soil cultivation on *Meloidogyne enterolobii* infection and on soil bacterial assemblages

**DOI:** 10.1101/2023.01.27.525929

**Authors:** Josephine Pasche, Janete A. Brito, Gary E. Vallad, Jeremy Brawner, Samantha L. Snyder, Ellen A. Fleming, Jingya Yang, Willian C. Terra, Samuel J. Martins

## Abstract

Soil cultivation may change the soil microbiome and alter interactions between plants and parasites. The objective of this work was to evaluate temporal changes in plant health, microbiome abundance, bacterial diversity and the plant-parasitic nematode, *Meloidogyne enterolobii* incidence in two soil fields with different agricultural uses. Soil samples were collected from a commercial tomato production field (agricultural soil) and a single-cultivation strawberry field (native soil). Samples for the second experiment were collected from the same fields the following year. Tomato plants cv. Yearly Girl were grown in a greenhouse and inoculated with *M. enterolobii*. After 45 days, plants were evaluated for the plant growth parameters, nematode reproduction, and soil bacterial assemblages were assessed using cultivation-independent sequencing methods (V3/V4 region of the rRNA 16S). Overall the average of fruit fresh weight in the second experiment was 2.4-fold to 14-fold higher than the first experiment. Moreover, there was a 80.5% decrease in eggs present per root system from the first experiment to the second. The relative abundance of bacterial assemblages from Experiment 1 to Experiment 2 changed for most of the top phyla (eg. *Actinobacteria, Bacteroidetes*, and *Chloroflexi*) and genera (eg. *Bacillus, Streptomyces*, and *Flavisolibacter*) and there was no change in microbial diversity between the two experiments. This study suggests that soil management can lead to an overall decrease in nematode reproduction and better crop yield, as well as a shift in the overall bacterial assemblages.

**Graphical Abstract:** 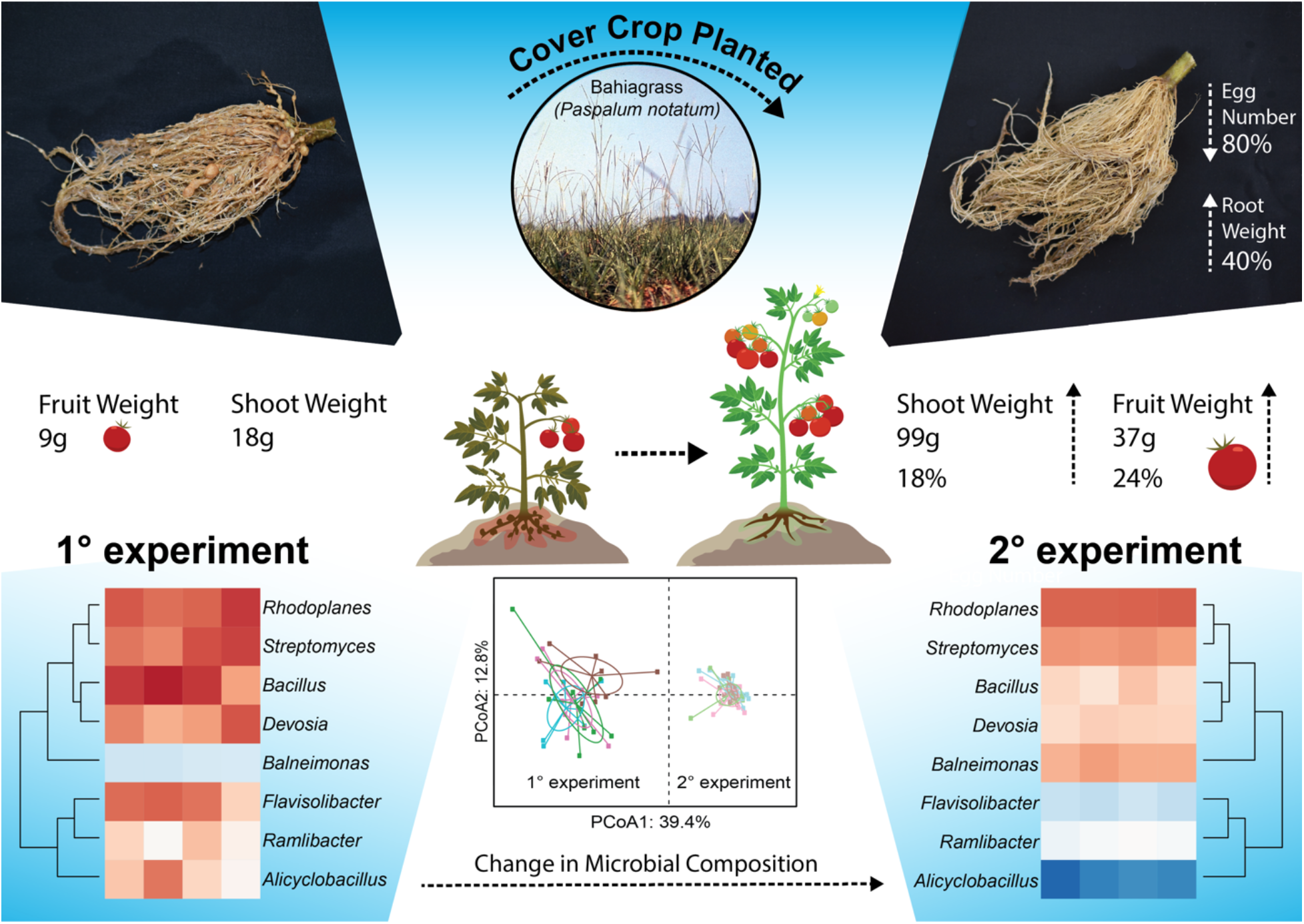

## Introduction

Root-knot nematodes, *Meloidogyne* spp., are sedentary endoparasites in many crops worldwide. *Meloidogyne enterolobii* (*M. enterolobii* hereafter) (Yang & Eisenback, 1983), in particular, is an emerging global threat to a wide range of plant hosts, causing chlorosis, stunting, root galls, and yield loss. *M. enterolobii* has earned the common name of “the guava root-knot nematode (GRKN)” due its ability to cause significant crop damage and even plant death on guava in South America (Carneiro et al. 2001). *M. enterolobii* is now considered a global threat to tomato (*Solanum lycopersicum* L.) production due to the lack of known resistance in commercially accepted varieties and yield loss of up to 65% (Centintas et al. 2007; Philbrik et al. 2020) and other crops such as guava and sweet potato (Khan et al. 2022; Schwarz et al. 2020). When *M. enterolobii* is established in the soil, it becomes difficult to manage the disease (Schwarz et al. 2020). Nematode infections can become even more problematic upon soil borne fungal infection, such as *Fusarium* spp., as a synergic effect between the two organisms may lead to greater nematode infections (Veloso et al. 2021; Khan et al. 2022). Moreover, due to its broad host range, high degree of virulence, and its ability to overcome root-knot nematode resistant genes in several important agricultural crops it is difficult to implement integrated pest management strategies to manage this plant-parasitic nematode. The application of highly toxic chemical nematicides, such as fumigants and non-fumigants, has been used to reduce the populations of plant-parasitic nematodes (PPNs) in soil. However, chemical products, especially nematicides, can be expensive and, in some cases, not economically viable to use. Additionally, nematicides are non-selective chemicals, acting on non-target organisms in the soil. In recent years, the application of certain highly toxic nematicides has been banned, which has limited nematode management options for growers (U.S. Environmental Protection Agency 2008; Zasada et al. 2010). Therefore, a more sustainable approach needs to be further developed to manage plant pathogenic nematodes, such as *M. enterolobii*.

Some biological control agents based on bacteria have recently been shown to be effective in controlling eggs, juveniles, and adults of *Meloidogyne* spp. (Stirling, 2014; Seid et al. 2015; Forghani and Hajihassani, 2020). In addition to the application of individual biocontrol agents to manage PPNs, changes in soil bacterial composition can affect *Meloidogyne* spp. populations. For instance, changes in soil conditions and agricultural practices can alter the bacterial composition of the soil, resulting in a reduction in the number of *Meloidogyne* spp. (Hermans et al. 2020; Madegwa et al. 2021). Similar to the changes caused by regular crop cultivation, the use of seasonal cover crops can also alter the soil microbiome (Kim et al. 2020). In this study, we hypothesized that successive crop cultivation will shift the microbial community in the soil and lead to a reduction in nematode reproductions.

The objective of this work was to assess the soil microbiome and plant health regarding infections caused by the emergent plant-parasitic nematode *Meloidogyne enterolobii* using soils collected from two fields with different agricultural uses.

## Material and methods

### Soil sample collection

Soil samples were collected at the Gulf Coast Research and Education Center, Wimauma, FL, USA field (39.6 m altitude, 27°45’23’’N and 82°13’29’’W). The soil type collected in this location is classified as fine sand according to the USDA Web Soil Survey. Two types of soil were collected: 1) located in a pasture/undisturbed soil with a limited production history, including cover crops (Ironclay peas and Buckwheat) for organic certification, a season of strawberry cultivation in the first year, followed by a Bahia grass (*Paspalum notatum*) cover crop in the second year (hereafter native soil); 2) located in a nearby field that had had successive tomato cultivation (agricultural soil) for the last 15 years. The soils were collected for the first and second experiments in 2021 and 2022, respectively, in the same areas and during the same period of the year (second week of June following harvest).

The soil was collected to a depth of 20 to 25 cm, with the first 2 cm of the soil surface layer being removed as its microbiome is highly influenced by aboveground changes. Soil was sampled from multiple areas in each field, and all soil samples were thoroughly mixed into a single composite. Before setting up the experiments, plant-parasitic nematodes were extracted from 200 cc of each soil type as described in the centrifuge floatation method (Cetintas et al. 2007). Nematode suspensions were examined using a light microscope (Olympus BH2), and checked for the presence of plant-parasitic nematodes, if any. Additionally, soil samples were sent out for physical chemical analysis at the Institute of Food and Agriculture Science Soil and Water Science Department of the University of Florida, Gainesville, FL.

The collected soils were sieved, homogenized, and used to fill 2 L pots. Pots remained under greenhouse conditions (temperature ca. 26°C, relative humidity ca. 67% and light intensity ca. 7,336 cd.sr m^−2^ s^−1^) at the University of Florida, in Gainesville, Florida, USA (54 m altitude, 29°63’90’’N and 82°35’60’’W). Plants were kept under greenhouse conditions as described previously and watered to field capacity.

### Nematode and Fungal inoculum preparation

*Meloidogyne enterolobii* isolate N01-000514 was obtained from Dade County and has been reared on tomato at the root-knot nematode collection, Division of Plant Industry, DPI/FDACS, Gainesville, FL, USA. Nematode eggs were extracted from infected roots using the Hussey and Barker (1973) method modified by Brito et al. (2020). Briefly, eggs were extracted from the roots using 1% NaOCl. The NaOCl solution was added until the liquid just fully covered the roots. The root-NaOCl mixture was blended on the highest setting for approximately 30 seconds, and the blended remains were added to a series of sieves (No.200 and No.500 Mesh, respectively). The nematode suspension was rinsed with a continual stream of tap water until there was no visible trace of bleach on the sieves. The eggs caught by the No.500 sieve were then resuspended in water to a final concentration of 1,000 eggs mL^−1^.

*Fusarium oxysporum* f.sp. *lycopersici* race 3 (*F.o.l*) was obtained from Dr. Vallad’s Lab located at the Gulf Coast Research and Education Center, Florida. The fungus was cultured in LB medium at 25°C for 4 days. To prepare the spore suspension, cheesecloth was used to separate fungal hyphae and spores. The concentration of spore suspension was calculated with a hemocytometer and adjusted to 10^5^ spores mL^−1^ for each assay.

### Plant inoculation methods

Three groups of inoculums were used in the experiment: *F.o.l*. only, *M. enterolobii* only, and *F.o.l*. x *M. enterolobii*, with sterile water used as a control. Four-week-old *S. lycopersicum* cultivar Early Girl plants (susceptible to *M. enterolobii*) were used in both assays. For the nematode inoculation, a suspension of *M. enterolobii* eggs at a volume of 5 mL (n=5000 eggs per pot) was added in each pot. Using a 15 cm falcon tube, 4 holes of ~1.5 cm wide and 6 cm deep were made in the form of a cross in the soil around the tomato root system. A 5 mL nematode solution was poured evenly into the holes, and immediately after inoculation, the holes were covered with soil. For the *Fusarium* inoculation, a volume of 5 mL of fungal spore suspension was drained into the soil near the base of the stem of each pot. Non-inoculated plants were used as the control. At 45 days after inoculation (dai), all plants were recovered and assessed for plant growth parameters (fresh weights) and nematode reproduction.

### Egg extraction

Eggs were extracted from the root systems of each experiment using the 1% NaOCl as described above.

### Soil DNA extraction and 16S sequencing

The soil microbiome DNA was extracted using 200 mg of soil from each pot and the Zymo Quick-DNA™ Fecal/Soil Microbe Miniprep Kit (Zymo Research, Irvine, CA). DNA purity was assessed using spectrophotometry (NanoDrop model, ThermoFisher, Waltham, MA), and DNA concentration was assessed by a Qubit 4 fluorometer (ThermoFisher, Waltham, MA). DNA was processed in the 16S (V3 and V4) region at the UF Interdisciplinary Center for Biotechnology Research (ICBR) using paired-end read sequencing on the Illumina MiSeq (2×300 bp) system.

### Experimental design and statistical analysis

Randomized complete block design was used for the *in vivo* tests with 7 reps per soil treatment. Data were submitted to analysis of variance (ANOVA) respectively for experiments 1 and 2. Kruskal-Wallis one way analysis of variance on Ranks (*P*<0.05) were applied for significant means for the two experiments. Tukey’s multiple range test was applied for comparison among biomass parameters: fresh fruit weight, fresh shoot weight, and root fresh weight. For all analyses, the assumption of normality was checked by Shapiro-Wilk and Kolmogorov-Smirnov tests prior to analysis. SigmaPlot® version 14.5 was used for statistical analyses, and SHAMAN was used for metataxonomic analysis from raw reads to statistical analysis (Volant et al. 2020). Sequences were clustered by operational taxonomic unit (OTU) at 97% similarity, and OTU taxonomy was assigned using the greengenes database (McDonald et al. 2012). For principal coordinates analysis (PCoA), an analysis of variance using the Bray-Curtis distance metric in a permutational multivariate ANOVA [PERMANOVA] test was conducted (P<0.05). Raw read data were submitted to the NCBI SRA under the accession number PRJNA914735.

## Results

Prior to beginning the experiment both soils were assessed for the presence of *Meloidogyne enterolobii* and other plant-parasitic nematodes through the Centrifugal Flotation method (Hussey and Barker, 1973), and no presence of any the plant-parasitic nematode was found in the soils. Additionally, the soils were assessed for their physical-chemical parameters, and the results are presented in Table 1.

**Table 1.**
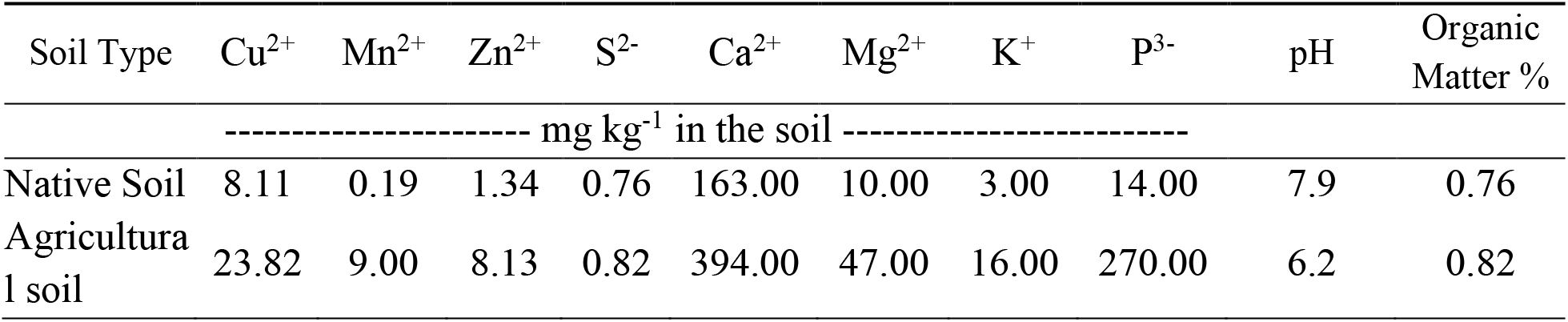
Soil chemical parameters assessment (organic matter content, macro- and micronutrients) for the two types of soils used in this study.

| Soil Type | Cu<sup>2+</sup> | Mn<sup>2+</sup> | Zn<sup>2+</sup> | S<sup>2-</sup> | Ca<sup>2+</sup> | Mg<sup>2+</sup> | K<sup>+</sup> | P<sup>3-</sup> | pH | Organic Matter % |
| --- | --- | --- | --- | --- | --- | --- | --- | --- | --- | --- |
|  | ----- mg kg<sup>-1</sup> in the soil ----- |  |  |  |  |  |  |  |  |  |
| Native Soil | 8.11 | 0.19 | 1.34 | 0.76 | 163.00 | 10.00 | 3.00 | 14.00 | 7.9 | 0.76 |
| Agricultural soil | 23.82 | 9.00 | 8.13 | 0.82 | 394.00 | 47.00 | 16.00 | 270.00 | 6.2 | 0.82 |

### Nematode and fungal results

A difference between the two experiments was found regarding the number of eggs of *M. enterolobii*, with an 80.5% reduction in the number of eggs for experiment 2 compared to experiment 1 (P<0.001) (Figure 1).

**Fig. 1.**
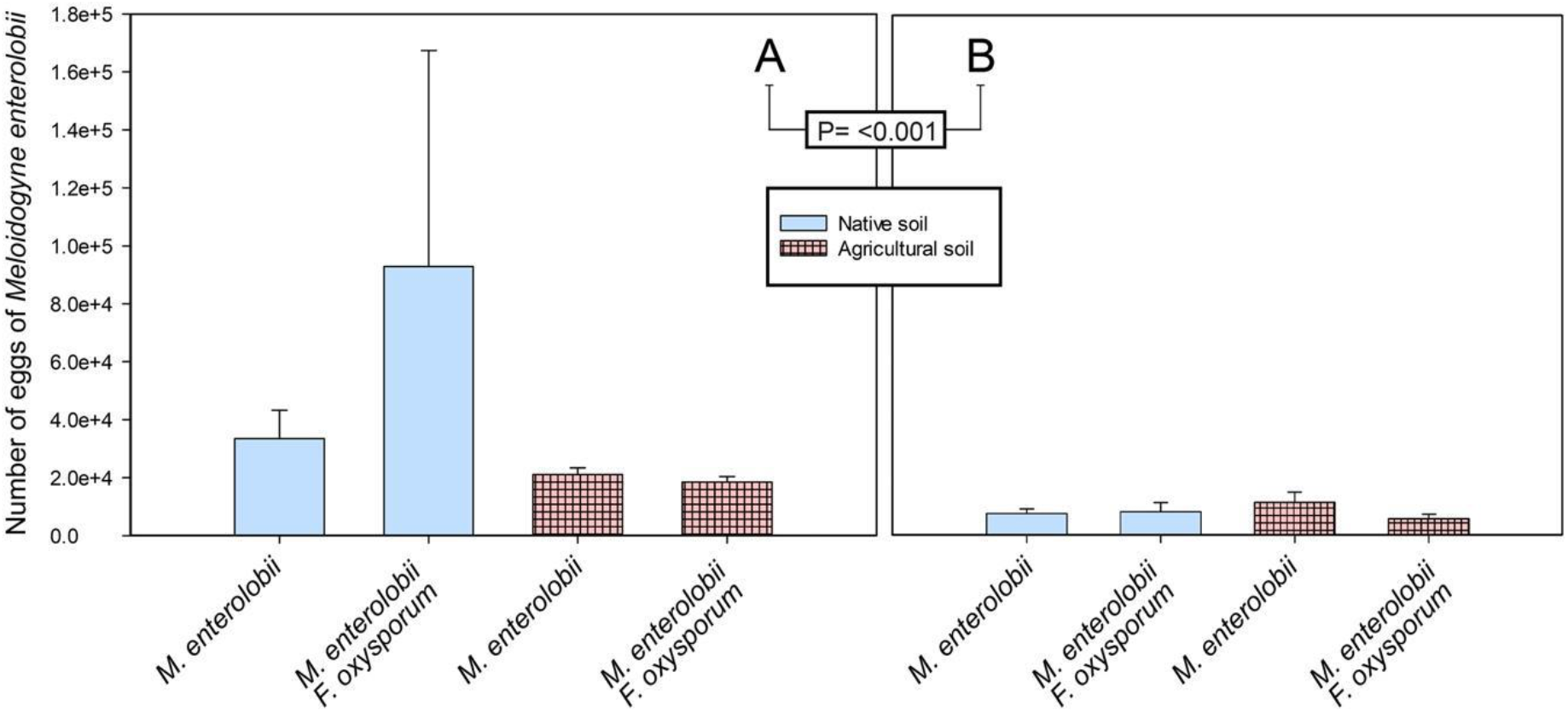
Average number of eggs of *Meloidogyne enterolobii* in roots of tomato plants cv. Yearly Girl at 45 days after egg inoculation and cultivation in greenhouse with soils collected in summer of 2021 (A) and summer of 2022 (B). Plants were infected either with the nematode only (5,000 eggs) or co-infected with spores of *Fusarium oxysporum* f.sp. *lycopersici* race 3 (10^5^ spores mL^−1^). Error bars represent ±SE.

On the other hand, no difference was found between the soils for both experiments 1 (P=0.477) and 2 (P=0.872) or among the treatments for both experiments 1 (P=0.572) and 2 (P=0.648) regarding the number of eggs.

### Plant biomass parameters

No difference was found for the two types of soils in the first experiment regarding fruit fresh weight (ffw), shoot fresh weight (sfw), and root fresh weight (rfw) (P=0.425, P=0.138, and P=0.085, respectively) or in the second experiment for ffw, sfw, and rfw (P=0.100, P=0.226, and P=0.631, respectively). On the other hand, there was a statistical difference when the experiments were compared for fresh fruit weight, fresh shoot weight, and root fresh weight (P=0.003, P<0.001, P<0.001, respectively), with a bigger yield and larger plant biomass for the second experiment compared to the first experiment. For instance, in the second experiment, the yield ffw was higher for all the treatments compared to the first experiment (Table 2).

**Table 2.** Tomato plants cv. Yearly Girl biomass parameters fresh fruit weight (ffw), fresh shoot weight (fsw), and root fresh weight (rfw) at 45 days after *Meloidogyne enterolobii* inoculation (5,000 eggs per pot) and/or with *Fusarium oxysporum* f.sp. *lycopersici (F.o.l*) race 3 (10^5^ spores mL^−1^) in greenhouse with soils collected in summer of 2021 (Experiment 1) and summer of 2022 (Experiment 2). Means of twelve replicates for each experiment. Same letters associated with the means are similar at the 5% level according to Tukey’s multiple range tests (columns). ±SE represents the standard errors.

| Treatment | Experiment 1*** |  |  |  |  |  | Experiment 2 |  |  |  |  |  |
| --- | --- | --- | --- | --- | --- | --- | --- | --- | --- | --- | --- | --- |
| | ffw | $\pm\text{SE}$ | sfw | $\pm\text{SE}$ | rfw | $\pm\text{SE}$ | ffw | $\pm\text{SE}$ | sfw | $\pm\text{SE}$ | rfw | $\pm\text{SE}$ |
| Control | 14.22a | 1.70 | 21.48a | 1.52 | 7.86b | 0.72 | 34.35ns | 11.21 | 89.30ns | 11.99 | 16.47b | 1.49 |
| *F.o.l.* | 15.16a | 2.77 | 23.34a | 2.26 | 9.34ab | 1.09 | 52.97ns | 11.30 | 123.86ns | 14.43 | 26.36a | 4.70 |
| *F.o.l.* + *M. enterolobii* | 2.35b | 1.56 | 12.64bc | 1.37 | 12.73a | 1.25 | 34.01ns | 9.48 | 96.90ns | 8.66 | 25.10b | 3.58 |
| *M. enterolobii* | 3.37b | 1.76 | 14.15b | 2.09 | 12.41a | 0.87 | 24.55ns | 10.83 | 86.27ns | 11.41 | 29.08a | 3.37 |

\*\*\*Significant at the 0.001 probability level (Experiment VS Experiment 2) ns = not significant

The increase in ffw in the second experiment varied from 2.4-fold to 14-fold, respectively, for the control treatment and when plants were co-inoculated with *M. enterolobii* and *F.o.l*. Additionally, there was a decrease in shoot fresh weight (sfw) when plants were inoculated with *M. enterolobii* alone (34%) or with *F.o.l*. (41%) compared to the control in the first experiment. However, in the second experiment there was no difference among the treatments regarding sfw (P=0.087). Unexpectedly, for both the first and second experiments, a reduction in the root fresh weight (rfw) was found in the control compared to plants infected with *M. enterolobii* alone or with *F.o.l*. (first experiment), and plants infected with *F.o.l*. alone or *M. enterolobii* alone (second experiment). In the second experiment, overall root fresh weight (rfw) increased (40%), along with shoot fresh weight (18%) and fruit fresh weight (24%).

### Microbiome results

The microbiome profiles of experiment 1 and experiment 2 showed statistically significant separation in diversity (permutational multivariate ANOVA [PERMANOVA], *P*<0.001; Figure 2A).

**Fig. 2.**
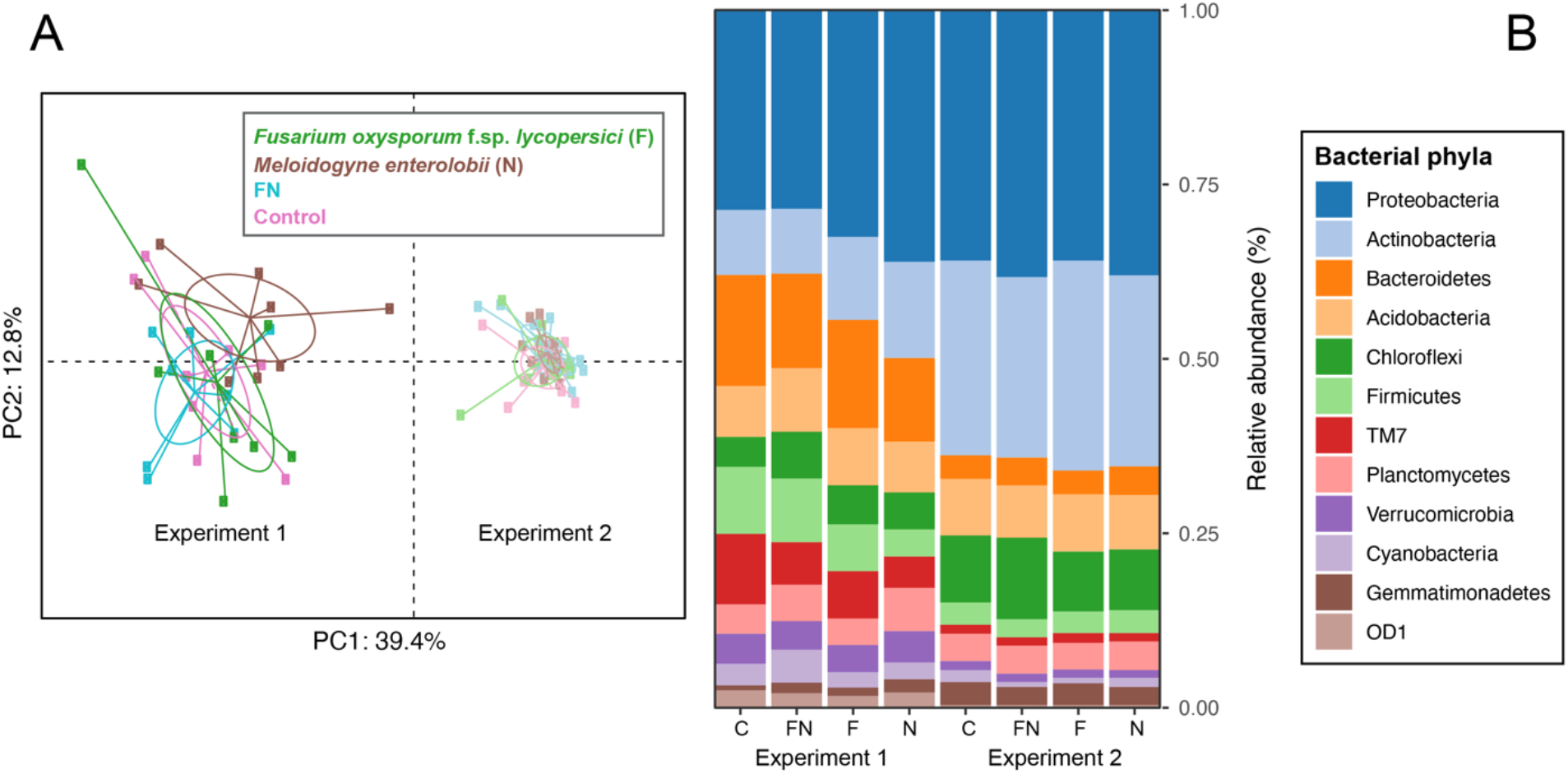
(A) Principal coordinates analysis (PCoA) based on Bray-Curtis distance metric. Different colors represent different treatments, and lighter colors represent data from experiment 2. The percentage of the variation explained by the plotted principal coordinates is indicated on the axes. (B) Relative abundance of operational taxonomic units (OTUs) of the twelve most abundant bacteria at the phylum level. C=control, FN = *Fusarium oxysporum* f.sp. *lycopersici (F.o.l*) race 3 & *Meloidogyne enterolobii*; F = *F.o.l.*; N = *M. enterolobii*.

Additionally, the separation between experiments is better explained by the highest Principal Coordinate 1 (PC1 = 39.4%). There was a shift in the bacterial community structure for the twelve most abundant bacterial phyla from experiment 1 to experiment 2 (Figure 2B). For both experiments, the most abundant phylum was Proteobacteria. The phyla Bacteroidetes, Firmicutes, TM7, Verrucomicrobia, and OD1 were more abundant in experiment 1 than in experiment 2. In experiment 2, Actinobacteria, Chloroflexi, and Gemmatimonadetes were more abundant than in experiment 1.

While there is a change in relative abundance of bacterial assemblages from experiment 1 to experiment 2, the Shannon Diversity Index showed no change in diversity between the two experiments (Figure 3A).

**Fig. 3.**
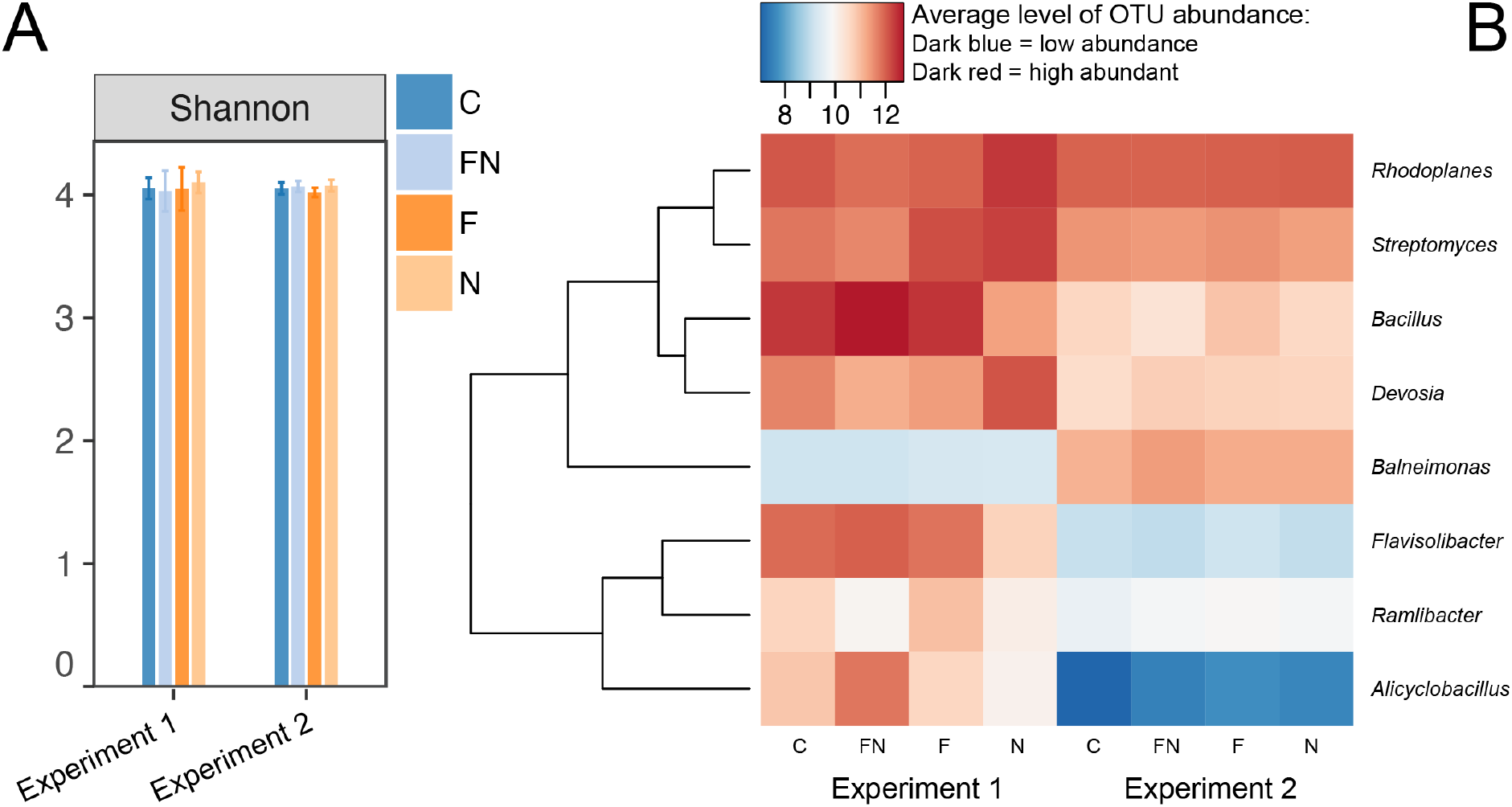
Comparison between experiment 1 and experiment 2 on: Shannon diversity (A) Heatmap of the average level of OTU abundance of the top 8 most contrasting bacteria at the genus level (B). C=control, FN = *Fusarium oxysporum* f.sp. *lycopersici (F.o.l*) race 3 & *Meloidogyne enterolobii*; F = *F.o.l*.; N = *M. enterolobii*.

The average level of OTU abundance shifted in the majority of the top 8 most abundant bacterial genera from experiment 1 to experiment 2. The genera in experiment 2 showed an overall decrease in abundance except for *Balneimonas*, which increased in abundance from experiment 1. The most abundant genus was *Bacillus*, in which the highest level of OTU abundance was found in experiment 1 under the FN treatment. This group corresponded to the highest infection rate based on the number of eggs present (Figure 1A). In experiment 2, the genus with the highest abundance was *Rhodoplanes*.

## Discussion

In this study we showed that soil with different uses can lead to an overall decrease in nematode infection (Figure 1) as well as a shift in the overall bacterial assemblages (Figure 2). Raaijmakers and Mazzola (2016) conducted a study focusing on successive host plant cultivation and pathogen reinfection and its effect on disease severity over time. The authors found that microbiome composition changed and increased in diversity with successive host plant cultivation. In our study, we found that soils collected from the first and second experiment from two different fields led to a different microbiome relative abundance. Unlike Raaijmakers and Mazzola’s (2016) study, the bacterial diversity did not change between experiments for our study.

In the first experiment, the overall nematode infection was higher in native soil but decreased in the second experiment, as the soil was more utilized. Previous studies have shown that more intensive use of the soil leads to more microbial abundance and a decrease in the nematode community. A study conducted by Yang et al. (2021) assessed the impacts of differing types of land use on nematode populations, and they found that more intensive use of the soil had negative effects on soil nematodes. A similar study by Hirschfeld et al. (2020) also found that when a native soil was shifted to an agricultural soil, the overall nematode population decreased. Furthermore, Knight et al. (2013) tested twelve soils from city-owned vacant lots, which could potentially be used in urban agriculture, and verified that PPN were among the most influential biological parameters for soil health.

The co-inoculation of *M. enterolobii* and *F.o.l*. presented a stronger reduction of plant shoot weight in the first experiment, which suggests that the interaction of *M. enterolobii* and *F.o.l*. was able to synergistically decrease plant biomass. Similar results were found by Wagner et al. (2022), who found that, in cotton, the interaction between *Fusarium* and *Meloidogyne* significantly reduced plant biomass as compared to each pathogen on their own. These results are supported by Regmi et al. (2022), who described the synergistic effect of a co-infection of *Fusarium* and *Meloidogyne* in tomato plants, which decreased overall plant biomass. Experiment 2 showed an overall drop in the nematode population compared to experiment 1. There is a possibility that the difference was influenced by the cover crop, Bahia grass (*Paspalum notatum*), that was planted in the native soil between the experiments. Previous literature has shown cover crops to have the ability to decrease the severity of nematode infection. Khan et al. (2022) sought to evaluate the effect of summer cover crops on the severity of *Meloidogyne enterolobii* infection. The authors found the use of non-host and poor-host cover crops in the summer significantly reduced future nematode populations.

A plant experiencing distress from a biological or abiotic stressor will release root exudates, or a chemical “cry for help,” to increase the presence of beneficial microorganisms that can minimize damage caused by the stressor (Rizaludin et al. 2021). In experiment 1, the genus found with the highest overall abundance was *Bacillus*, specifically in the group with the addition of both pathogens, *Fusarium* and *M. enterolobii* (Figure 3B). Previous studies have addressed *Bacillus* as a biocontrol agent for the soilborne pathogens *Fusarium* and *M. enterolobii* (Khan et al. 2018; Vieira de Carvalho Júnior et al. 2022). Specifically, Singh et al. (2021) conducted a study in which they assessed the biocontrol potential of *Bacillus subtilis* against *Meloidogyne incognita* and *Fusarium oxysporum*. The authors found that there is potential for *B. subtilis* to act as a biocontrol agent against both soilborne pathogens.

We found a significant increase in root fresh weight with the presence of an *M. enterolobii* infection (Table 2). A study done by Almeida et al. (2022) analyzed tomato root growth under differing conditions, including the addition of *Meloidogyne enterolobii*, to a commercial substrate. The authors found the root fresh weight to be significantly higher with the addition of a GRKN infection. Increased root growth can often be accompanied by an increase in feeding sites and galls (Sohrabi et al. 2020). However, other studies have shown a decrease in root weight with an increase in nematode galls (Ganeshan et al. 2019; Karabörklü et al. 2022).

In conclusion, this study verified that anthropogenic changes in the soil (eg. crop cultivation) can lead to major changes in the soil microbiome between experiment 1 and experiment 2, favor some bacteria over others, and ultimately lead to less nematode infection in tomato plants caused by the emergent plant parasitic nematode *Meloidogyne enterolobii*. Further studies could focus on specific soil manipulations (eg. soil amendments) in native soil as a way to build soil health to become suppressive against plant parasitic nematodes.

## Acknowledgement

This work was supported by the USDA National Institute of Food and Agriculture (NIFA), Hatch project N° 1024881 and the NIFA grant project N° 2021-68013-33758. The authors are also thankful to Beatriz Franceschi for her help with soil collection and Malia M. Fortney and Dr. Diego A. H. S. Leitão and Ruimin Xue for their help with the nematode extraction assay.

